# Extracting electromyographic signals from multi-channel local-field-potentials using independent component analysis without direct muscular recording

**DOI:** 10.1101/2022.11.15.516633

**Authors:** Hisayuki Osanai, Jun Yamamoto, Takashi Kitamura

## Abstract

Electromyography (EMG) has been commonly used for precise identification of animal behavior. However, they are often not recorded together with in vivo electrophysiology due to the need for additional surgeries and setups and the high risk of mechanical wire disconnection. While independent component analysis (ICA) has been used to reduce noise from field potential data, there has been no attempt to proactively use the removed “noise”, of which EMG signals are thought to be one of the major sources. Here, we demonstrate that EMG signals can be reconstructed without direct EMG recording, using the “noise” ICA component from local field potentials. The extracted component is highly correlated with directly measured EMG, termed as IC-EMG. IC-EMG is useful for measuring an animal’s sleep/wake, freezing response and NREM/REM sleep states consistently with actual EMG. Our method has advantages in precise and long-term behavioral measurement in wide-ranged in vivo electrophysiology experiments.

**Highlights:**

- EMG signals can be extracted from LFP signals without direct muscular recording
- The extracted signal is highly correlated with direct EMG recording signals
- The extracted signal is useful in measuring animal behaviors as well as actual EMG
- This method contributes to precise and stable long-term behavior measurement

**In brief:** Osanai et al. demonstrate electromyography (EMG) signals can be extracted from multi-channel local field potential (LFP) recordings using blind-source-separation technique without direct measurement of muscle activity. The proposed method adds precise and long-term behavioral measurements with EMG information in wide-ranged in vivo electrophysiology experiments.

## Introduction

While neural correlates of animal behavior are examined most frequently with video observation of an animal’s movement, physiological signals such as electromyography (EMG) are also often used to monitor the small movements of the animal that can be missed in video observation. In neuroscience studies, EMG has been used to accurately assess sleep/wake states ^1–6^ and freezing behavior ^7–9^. However, despite the benefits of accurate behavior assessment, EMG is often not obtained together with brain activity recording because it requires additional surgery and setups ^7, 10–13^. Furthermore, in the case of implanting a silicon probe, most commercial headstage preamplifiers are not designed with extra pins to obtain EMG signals, thus an additional preamplifier is required which increases total implant weight, potentially interfering with the normal behavior of small animals such as mice. Moreover, a high risk of EMG electrode wire breakage during experiments ^14–16^ hampers long-term experiments.

Field potentials have been shown to be correlated with an animals’ behaviors such as sleep/wake conditions ^1, 2, 17–20^ and fear freezing behavior ^8, 21, 22^. Field potential signals are often electrically referenced to one of the electrodes placed in a brain to suppress excessive common noise generated from the outside of the brain ^19, 23–25^. On the other hand, because the field potential signals can be distorted by the electrical signals at the reference electrode site^24–26^, an electric reference is frequently set outside of the brain, (e.g. a skull screw) ^27–29^. In this case, in order to minimize the noise arising from the distal reference electrode setting, multiple methods have been proposed such as common average reference ^30^, current source density analysis ^31–33^, and application of independent component analysis (ICA) ^34^.

ICA is a blind source separation technique which isolates temporally independent source signals ^35–38^. This technique has been well established for decades in “denoising” recorded neural signals by isolating and deleting the noise components from the field potential data, in which EMG artifacts are considered to be a major noise source ^34, 39–43^. However, although the putative EMG component isolated by ICA has been the target of noise suppression, it has not yet been investigated to what extent this component correlates with real EMG signals. Thus, the component has not been proactively used as a measurement of animals’ behavioral states.

Here, we hypothesized that ICA can reconstruct EMG signals from multichannel local field potential signals obtained by a silicon probe or tungsten wire electrodes without direct measurement of muscular signals. To investigate the correlation between independent component (IC) and EMG signals, we simultaneously implanted a silicon probe into a mouse’s hippocampus and wire electrodes into the neck muscles. Local field potentials (LFPs) were recorded referenced to a skull screw over the cerebellum. Next, we obtained ICs from the LFPs using ICA, and demonstrated that EMG-like high frequency components (IC-EMG) could always be obtained from the LFPs. These components were shown to highly correlate with directly measured EMGs. We further demonstrated this IC-EMG can be used to measure sleep/wake activity and freezing behavior. Finally, we demonstrated that IC-EMGs can also be obtained from four-channel multisite brain-areas LFPs obtained by tungsten wire electrodes.

## Results

### IC-EMG is highly correlated with real EMG signals

LFP signals were obtained from a 32-channel silicon probe implanted in the mouse hippocampus, using a skull screw for electric reference above the cerebellum (Figs. 1A-D). At the same time, EMG signals were obtained from the left and right neck muscles (Fig. 1C, E). The ICs were obtained by ICA from the recorded multichannel LFP signals (Fig. 2A). Figure 2Ab illustrates one of the obtained IC (IC#1) showing EMG-like higher frequency activity than other ICs. The back-projected reconstructed signals on the original channels using this IC showed that the signals distributed uniformly along the channels (Fig. 2Ba). Deleting this IC clarified the weaker and slower activities, which had been previously contaminated with the noise-like large high-frequency activity in the original waveforms (Fig. 2Bb). Indeed, IC#1 showed a clear peak between 100-200 Hz in the estimated power spectrum density (Fig. 2C) and the weight distribution of this component was highly uniform compared to other ICs (Fig. 2D). In LFP data obtained from ten mice, only one IC showing uniform weight distribution (standard deviation (std) < 0.1) was consistently obtained from individual mice (Fig. 2Ea). To focus on uniformly distributed ICs, we classified ICs having a minimum std among ICs obtained in individual mice and a std < 0.1 as a uniform-IC. The uniform-IC also showed the maximum mean value of its weight distribution and tends to have large amplitude activities (Fig. 2Eb; uniform-IC vs. other ICs: weight mean = 0.97 ± 0.007 vs. 0.21 ± 0.013, t_(265)_ = 10.99, p = 2.15e-23; weight std = 0.019 ± 0.003 vs. 0.36 ± 0.009, t_(265)_ = −7.44, p = 1.42e-12; Amplitude ratio = 0.88 ± 0.090 vs. 0.20 ± 0.013, t_(265)_ = 9.88, p = 8.35e-20, respectively). In addition, the uniform-IC always had a peak frequency between 100-200 Hz similar to the real-EMG power spectrum density (Fig. 2F), indicating the uniform-IC reflected EMG signals.

**Fig. 1.**
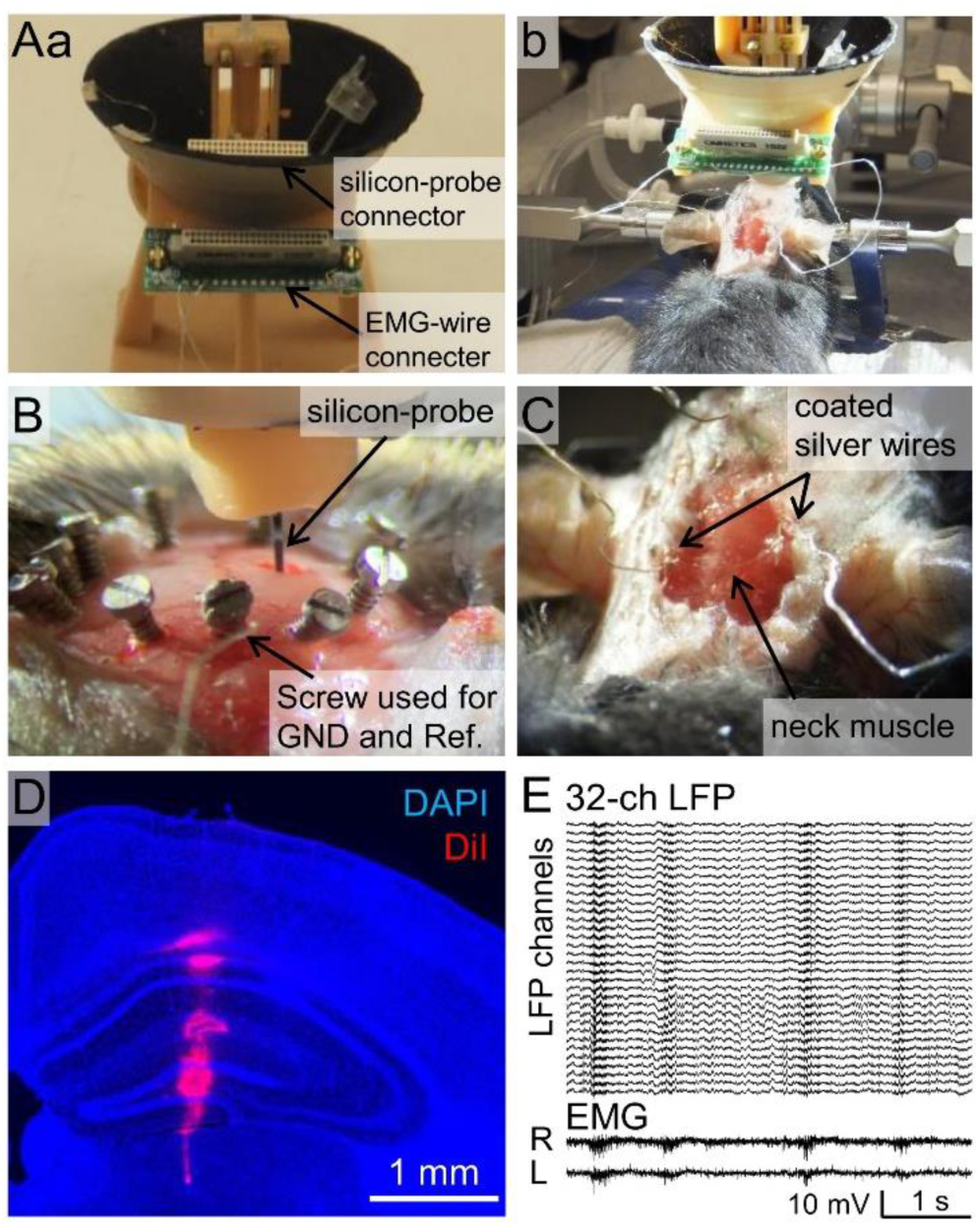
LFPs and EMG recording setups. (A) A microdrive system for implanting a silicon-probe and EMG electrode wires, placed on a jig (a) and during surgery (b). (B) Skull screw used for an animal’s ground and reference. The silicon probe was inserted into the brain in this picture. (C) EMG electrode wires placement. The tips of the coated wires were exposed and looped into the left and right neck muscles. (D) The trace of silicon probe implantation into HPC. The probe was coated with DiI before implantation. (E) Representative LFP traces from a 32-ch silicon probe referencing the skull screw (top), and right (R) and left (L) neck EMG traces (bottom).

**Fig. 2.**
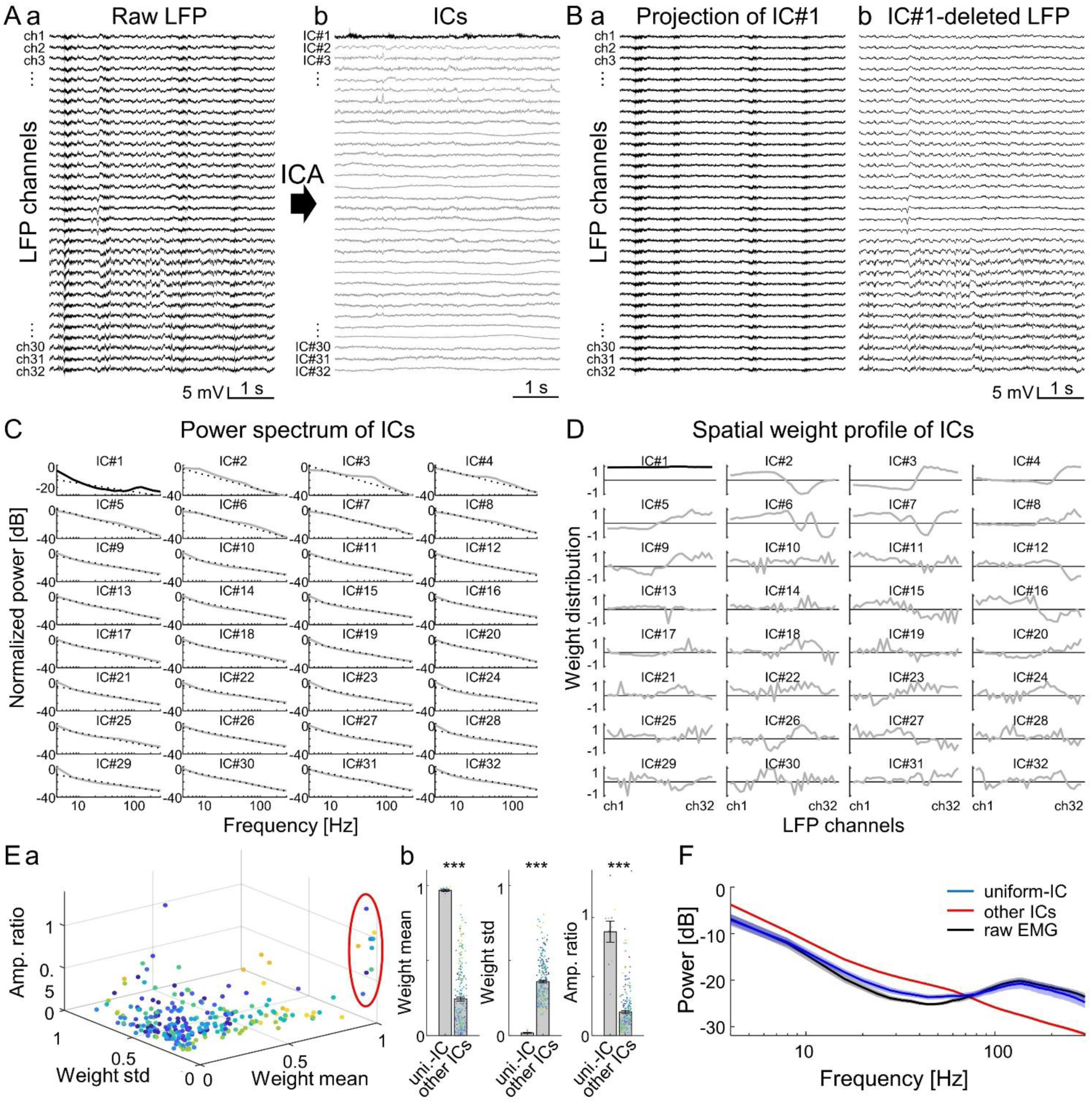
Identification of putative EMG component from LFP by ICA. (A) Raw LFP traces (a) and their independent components (ICs) separated by ICA (b). Note that IC#1 (black trace) showed higher frequency activity than other ICs (gray traces) similarly to the EMG traces in Fig. 1E. (B) (a) Projection of IC#1 to the original LFP channels. (b) The cleaned LFP obtained by deleting IC#1 from the raw LFP (b). The EMG-like high-frequency activities disappeared in IC#1-deleted LFP. (C) Log-log plots of the normalized power spectrum densities of each IC shown in *A*. IC#1 showed peak power at the range >100 Hz. Black dotted lines indicate the power of aperiodic exponent (see Methods). (D) The weight distribution of ICs shown in *A*. IC#1 had the most-uniform (minimum standard deviation (std)) distribution. (E) (a) Scatter plot of IC’s mean weight, std, and amplitude (see Methods), obtained from ten mice. Red circle indicates the ICs classified as uniform-IC in each mouse. (b) Averaged parameters of IC-EMGs and other ICs. Dots of each color indicate different mice. (F) Averaged normalized power spectral density of the uniform-IC (blue) other ICs (red), and raw EMG (black).

We next investigated to which extent the uniform-IC correlates with real-EMG signals. Figure 3A illustrates the filtered signals of uniform-IC and the real EMG, and Figure 3B shows the time course of their amplitude calculated by root-mean-square (RMS) with 100 ms time-windows. As we expected, the uniform-IC and the real-EMG signals were highly correlated, with correlation coefficients that were more than 0.9 (Fig. 3C and D).

**Fig. 3.**
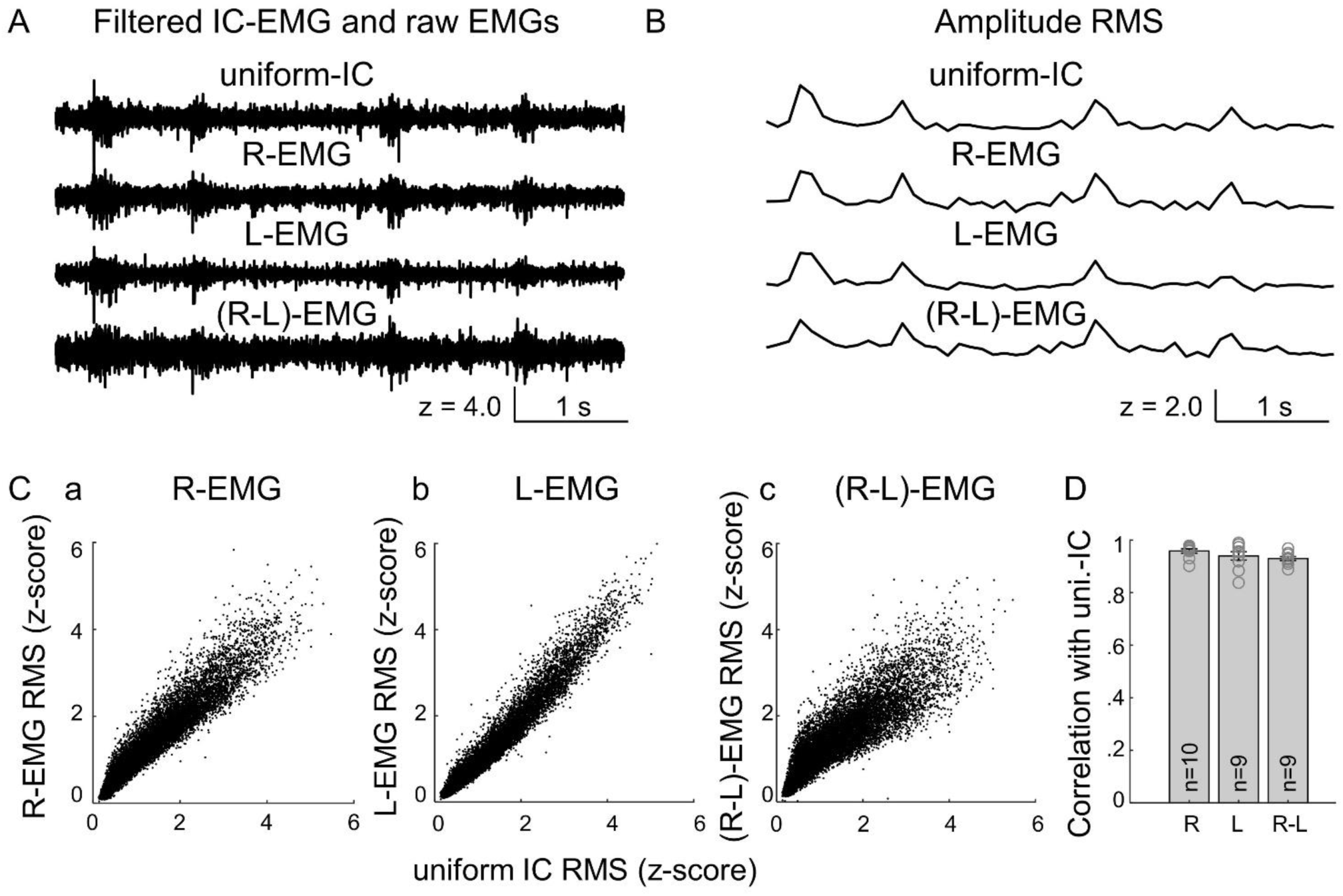
Correlation of uniform-IC and raw EMG signals. (A) 50-500Hz filtered waveforms of uniform-IC (IC#1 in Fig. 2), right-neck EMG (R-EMG), left-neck EMG (L-EMG) and, Δ(right-to-left) EMG ((R-L)-EMG) which is obtained by subtracting L-EMG from R-EMG to remove the effect of the common reference (skull-screw reference). (B) Root-mean-square (RMS) with 100 ms-bin time windows of each trace in A. (C) Correlation of uniform-IC with each EMG. (a) with R-EMG, (b) L-EMG, and (c) (R-L)-EMG. (R-L)-EMG is less correlated with IC-EMG than R- or L-EMG because the common reference effect is absent due to the subtraction, but still shows high correlation. (D) Average Pearson’s correlation coefficient of uniform-IC and real EMG from ten mice. L-EMG data from one mouse is missing due to the wire break during experiments. Correlation between uniform-IC and R-EMG = 0.96±0.008, L-EMG = 0.94±0.016, (R-L)-EMG = 0.93±0.007.

Therefore, EMG signals can be reconstructed from skull-screw-referenced multichannel LFP data by ICA without direct EMG recordings. These signals are represented by the uniform-IC, which hereafter we termed IC-EMG. The EMG-like components were not obtained from brain-referenced LFPs in most cases (eight out of ten mice), since EMG information was absent due to subtraction (Fig. S1A-C). We observed the EMG-like components from the brain-referenced LFPs only when the noise-level of LFPs was different between the recording channels, such that brain-referencing cannot completely reduce the noise (Fig. S1D-F, S2). Thus, brain-referenced LFP is not ideal for obtaining EMG signals using ICA.

### IC-EMG provides behavior measurements

Next, we examined whether the IC-EMG is useful for annotating animal’s behavioral states. Mice were placed into and allowed to move freely in the open field chamber (Fig. 4A). LFPs and EMGs were recorded while the mice were in the chamber (Fig. 4B), together with video tracking of animals’ positions. We categorized animal behavioral states from video into Sleep, Quiet Awake (QAW), and Active, classified according to the animals’ average head speed (Fig. 4C; see Methods). Amplitudes of IC-EMG and real-EMG were quantified in ten mice during each behavioral state. Fig. 4D shows that there were significant differences in the IC-EMG amplitudes, similarly to the real EMG, between Sleep, QAW, and Active behavioral states (Fig. 4D).

**Fig. 4.**
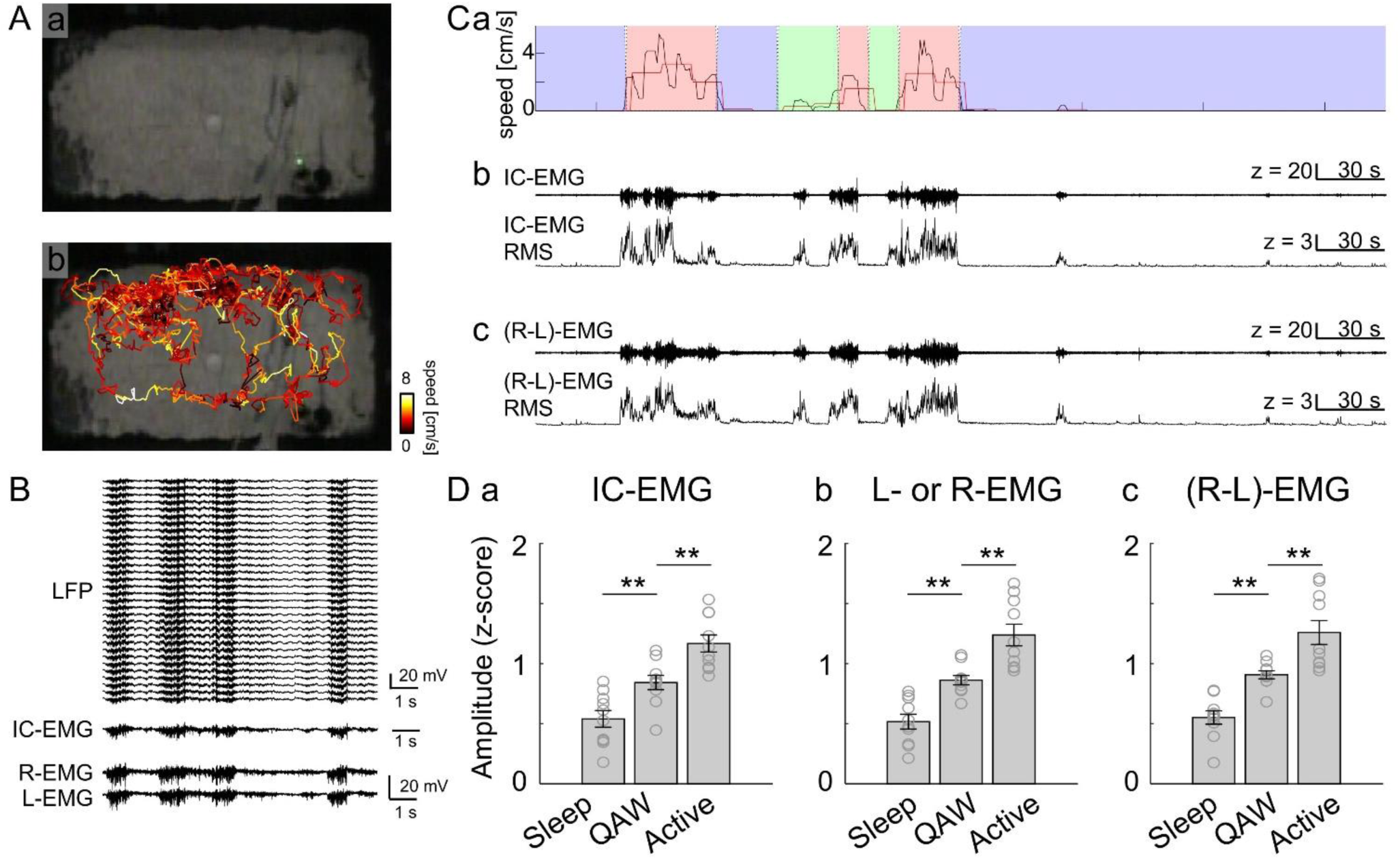
IC-EMG can provide measurements of animal behaviors. (A) The recording chamber (a) and head position tracing (b) during LFP and EMG recording. (B) Simultaneous recording of LFP and EMG when the mouse was freely moving in the recording chamber. EMG-IC was obtained from raw-LFP by ICA. (C) Speed of the animal’s head, traces of IC-EMG and raw EMG and their amplitudes at the same time. (a) head position speed (black line) and its average during each 15s epoch (red line), (b) z-scored IC-EMG and its RMS amplitude, (c) z-scored raw EMG and its RMS amplitude. Animal behavior was categorized into Sleep (blue), Quiet Awake (QAW, green), and Active (red) based on the video data. (D) Amplitude of IC-EMG (a, n = 10), L- or R-EMG (b, n = 10), and (R-L)-EMG (c, n = 9) during different animal behaviors. Amplitudes between Sleep vs. QAW vs. Active: IC-EMG: 0.54 ± 0.070 vs. 0.84 ± 0.060 vs. 1.17 ± 0.071, p = 0.010 (Sleep/QAW) and p = 0.005 (QAW/Active); L- or R-EMG: 0.52 ± 0.061 vs. 0.86 ± 0.039 vs. 1.24 ±0.090, p = 0.003 (Sleep/QAW) and p = 0.001 (QAW/Active); (R-L)-EMG: 0.55 ± 0.055 vs. 0.91 ± 0.033 vs. 1.26 ±0.100, p = 0.003 (Sleep/QAW) and p = 0.003 (QAW/Active).

Freezing behavior has been demonstrated as an indicator for fear response in both innate and conditioned reactions ^44–47^. We furthermore demonstrate that IC-EMG is useful for fear freezing detection, consistent with the previous studies showing that EMG can be used as a measure of fear freezing behavior ^7^. Mice were conditioned in the tone fear conditioning paradigm while their fear freezing was recorded with video observation, along with LFPs and real EMG signals (Fig. 5A). For conditioning, mice were exposed to three pairings of a 20 s tone and a 2 s foot shock on Day1. Then, fear freezing behaviors in response to tone were monitored on Day2 in a different context and scored by animal’s motion in the recorded video. At the same time, IC-EMG was obtained from the recorded LFPs, and the amplitudes of IC-EMG and real EMG were examined during animals showed freezing and non-freezing behavior (Fig. 5B). The mice actively explored the recording box before the tone delivery, while they showed increased freezing behaviors after the first tone delivery (Fig. 5B, C). From the LFP data of five mice, IC-EMG showed a significantly lower amplitude during the freezing state compared to the non-freezing state, similar to the results of real EMG signals (Fig. 5D). Together, these results demonstrated that IC-EMG is useful for annotating animal behavior such as sleep/quiet-awake/active behavioral states and freezing/non-freezing states.

**Fig. 5.**
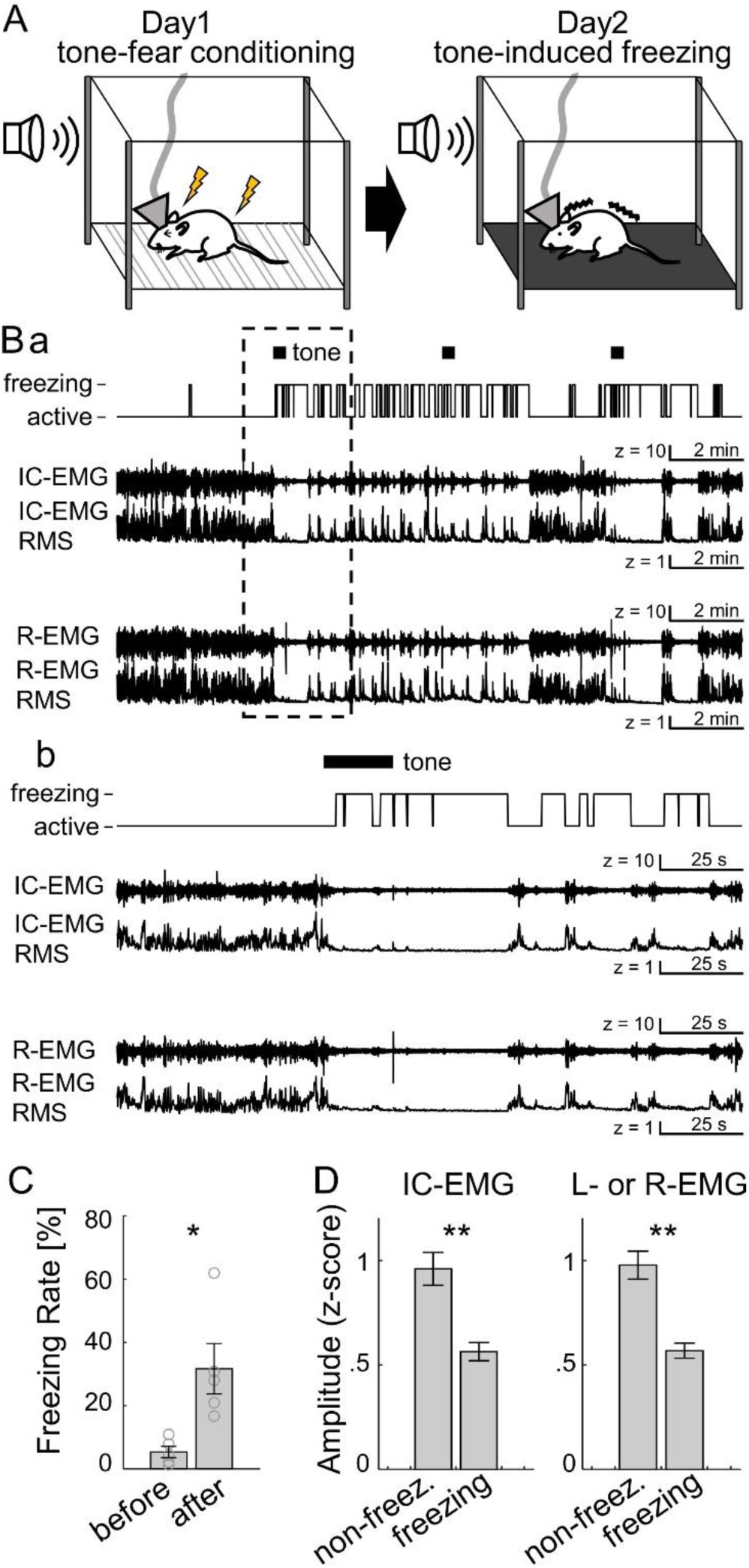
IC-EMG can be measurements of animal’s fear freezing. (A) Schema of the tone fear conditioning task. (B) Animals’ freezing behavior at day2, IC-EMG and raw EMG traces, and their RMS amplitudes during whole sessions (a) and 1 min before and after the tone onset (b). (C) Freezing rate during habituation session and test sessions (n = 5) before and after the first tone delivery. Freezing rate before vs. after the tone = 5.4 ± 1.8% vs. 31.7 ± 8.0%, respectively, t_(8)_ = −3.22, p = 0.012. (D) Amplitude of IC-EMG and L- or R-EMG during animal’s active/freezing states. (R-L)-EMG was not analyzed because one side of the EMG wires was broken during foot-shock delivery in three out of five mice. The z-scored amplitudes during non-freezing and freezing states: IC-EMG: 0.96 ± 0.079 vs. 0.56 ± 0.043, t_(8)_ = 3.65, p = 0.007; EMG: 0.98 ± 0.067 vs. 0.57 ± 0.036, t_(8)_ = 3.37, p = 0.001, respectively.

### IC-EMG can improve precise REM sleep detection

We examined whether the IC-EMG is useful for precise annotation of the rapid eye movement (REM) sleep stage. REM/non-REM (NREM) sleep periods are often classified by high/low theta-to-delta ratio of field potential when the animal shows stationary behavior ^48, 49^. However, because many factors can increase theta power during QAW ^8, 21, 22, 50–55^ (see Discussion), the combination only of video recording and field potential recording has difficulty in fine discrimination of QAW and REM sleep stats. The most established sleep scoring method uses a combination of field potential and EMG ^56–60^. In this paradigm, sleep/wake is classified by low/high amplitude EMG activities in addition to the animal’s movements in the recorded video. Here, we demonstrate IC- EMG can be used for sleep scoring similarly to real-EMG recording.

LFPs and EMGs of the mice were recorded in the open field chamber or their home cages (Fig. 6A). The animal’s behaviors were first annotated with the video tracking of animals’ positions (Fig. 6Aa). High theta- (6–9 Hz)-to-delta (0.1–4 Hz) ratio periods were observed during the mice were stationary (Fig. 6Ab, c). In order to separate REM sleep periods from QAW state, the animal behaviors were further classified using IC-EMG information that the stationary periods with high IC-EMG amplitudes were re-categorized as being in the awake-state (QAW) (Fig. 6Ad), similarly to the use of the EMG (Fig. 6Ae). Then, REM/NREM/QAW states were annotated with the combination of the theta-delta ratio and the animal’s speed, or the combination of the theta-delta ratio, the animal’s speed, and the IC-EMG or the EMG (Fig. 6B). Some periods annotated as REM based only on the video (Fig. 6Ba) were categorized into QAW when the annotation was based on IC-EMG (Fig. 6Bb), similarly to the result based on the EMG (Fig. 6Bc), because of their large amplitudes of muscular activities during the periods which are not likely to occur during sleeping states. The dynamics of IC-EMG and EMG during IC-EMG- and EMG-based categorized REM periods were consistent with previous studies where EMG shows silent and occasional twitch activities ^61, 62^, and that body-movement occurs at the end of REM ^63^. To evaluate the extent to which the quality of the video-based and the IC-EMG-based state classification is close to the EMG-based method, the REM sleep periods were detected from six silicon-probe implanted mice (Fig. 6C). Because the sleep periods were more strictly annotated, the IC-EMG based method showed less false-positive rates of REM detection compared to the video-only based method (Fig. 6Da; video: 1.18 ± 0.38%, t_(5)_ = 3.13, p = 0.025; IC-EMG: 0.39 ± 0.20%, t_(5)_ = 1.98, p = 0.104, one-sample t-test with zero), and negligible levels of false-negative rates in IC-EMG (0.16 ± 0.12%, t_(5)_ = 1.27, p = 0.260, one-sample t-test with zero). Also, the IC-EMG-based REM period detection was similar (Cohen’s kappa ^64–66^) to the EMG-based method while the similarity of the video-only based method was less (Fig. 6Db; video: 0.91 ± 0.03, t_(5)_ = −2.89, p = 0.035; IC-EMG: 0.96 ± 0.02, t_(5)_ = −1.93, p = 0.112, one-sample t-test with kappa = 1). Therefore, the use of IC-EMG improves precise REM sleep detection similarly to the use of real-EMG by decreasing false-positive REM periods.

**Fig. 6.**
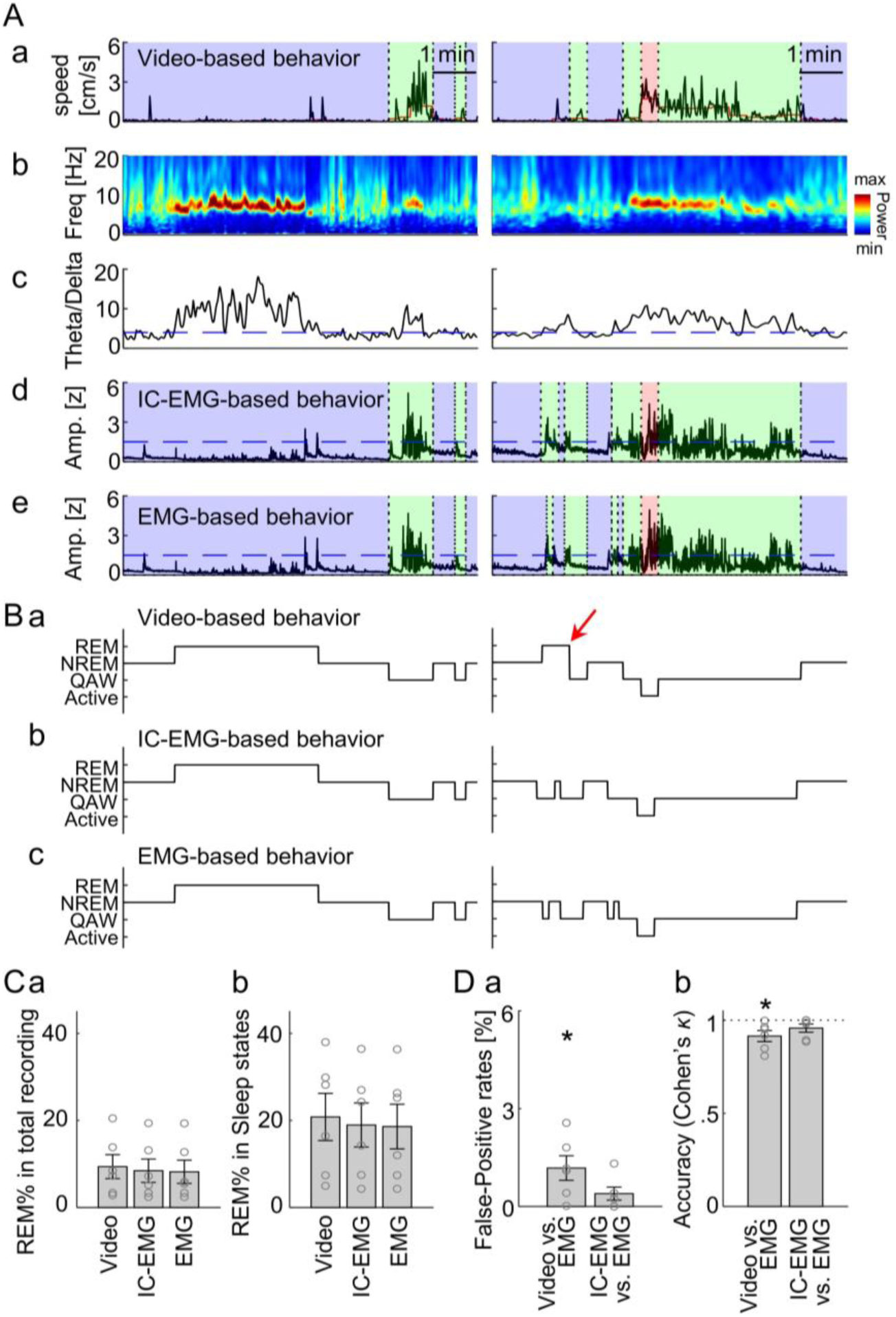
IC-EMG is useful for precise annotation of REM sleep scoring. (A) Sleep/wake annotation based on the video recording, IC-EMG and EMG. (a) head position speed (black line) and its average during each 15s epoch (red line). Animal behavior was categorized into Sleep (blue), QAW (green), and Active (red) based on the head speed. (b) Spectrogram of LFP at st. lm layer of CA1. (c) Trace of theta (6–9 Hz)-to-delta (0.1–4 Hz) ratio. Blue dotted line indicates ratio = 4. (d, e) Sleep states categorized based on video recording (a) were re-annotated into QAW if the amplitudes of the z-scored IC-EMG (d) and (R-L)-EMG (e) are high (see Methods). Black lines indicate RMS amplitudes of IC-EMG/(R-L)-EMG. Blue dotted lines indicate z = 1.5. (B) Sleep scoring based on the video recording (a), IC-EMG (b) and (R-L)-EMG (c). The states were classified into REM when the theta-to-delta ratio is ≥ 4 during Sleep state (see Methods). Red arrow indicates the REM period which is categorized based on the video was re-categorized as wake by IC-EMG- and EMG-based annotation due to their high amplitudes. (C) REM sleep duration from the six mice in the total recording time (a) (percentages based on video vs. IC-EMG vs. EMG: 9.38 ± 2.75 vs. 8.44 ± 2.68 vs. 8.21 ± 2.67), and in total sleep periods (b) (percentages based on video vs. IC-EMG vs. EMG: 20.80 ± 5.42 vs. 18.95 ± 5.06 vs. 18.59 ± 5.13). These amounts are consistent with the previous reports ^102, 103^. (D) The agreement of REM sleep scoring based on video and IC-EMG versus the scoring based on EMG; False-positive rates (a), and Cohen’s kappa values (b).

### IC-EMG can be obtained from multisite brain-area recording

Finally, we examined whether IC-EMG can be obtained from multisite brain-area recording data. We have previously recorded four channel LFPs in ACC, BLA, white matter of dorsal CA1, and ventral CA1 of mice ^67^. We had obtained the LFP signals referenced to the skull screw over the cerebellum, and ICs were separated by ICA from the recorded LFPs into ICs (Figs. 7A-D). Similar to the case of silicon-probe recordings (Fig. 2), one of the ICs consistently present in each of the four mice exhibited a highly uniform weight distribution (Fig. 7E, G). These components also showed a peak frequency between 100-200 Hz (Fig. 7F, H), which is consistent with the results of silicon-probe recordings (Fig. 7F), indicating a strong correlation with actual EMG signals. We examined the amplitude of IC-EMG when the mice explored the open field chamber. Again, consistent with the results of silicon-probe recordings, IC-EMG amplitudes were significantly higher during awake states than sleeping states (Fig. 7I). We finally examined IC-EMG amplitudes during freezing/non-freezing behaviors during the observational fear ^68–70^. The recorded mouse, as an observer, and his cagemate, as a demonstrator, were placed into the two chambers separated by a transparent plexiglass partition. On the recording day, the foot shock was delivered to the cagemate demonstrator which produced the observational fear response ^67^ (Fig. 7Ja). During the freezing period, IC-EMG of 4-channel LFP showed significantly lower amplitudes compared to the non-freezing state in the observer (Fig. 7Jb). Therefore, IC-EMG can also be obtained from multisite brain-areas LFP recording, and it provides behavior measurement of sleep/awake states and fear-freezing/non-freezing states. Moreover, these results also demonstrate that our method is able to reconstruct EMG signals from data obtained in the past, so long as an electrical reference was set on the animal’s skull screw.

**Fig. 7.**
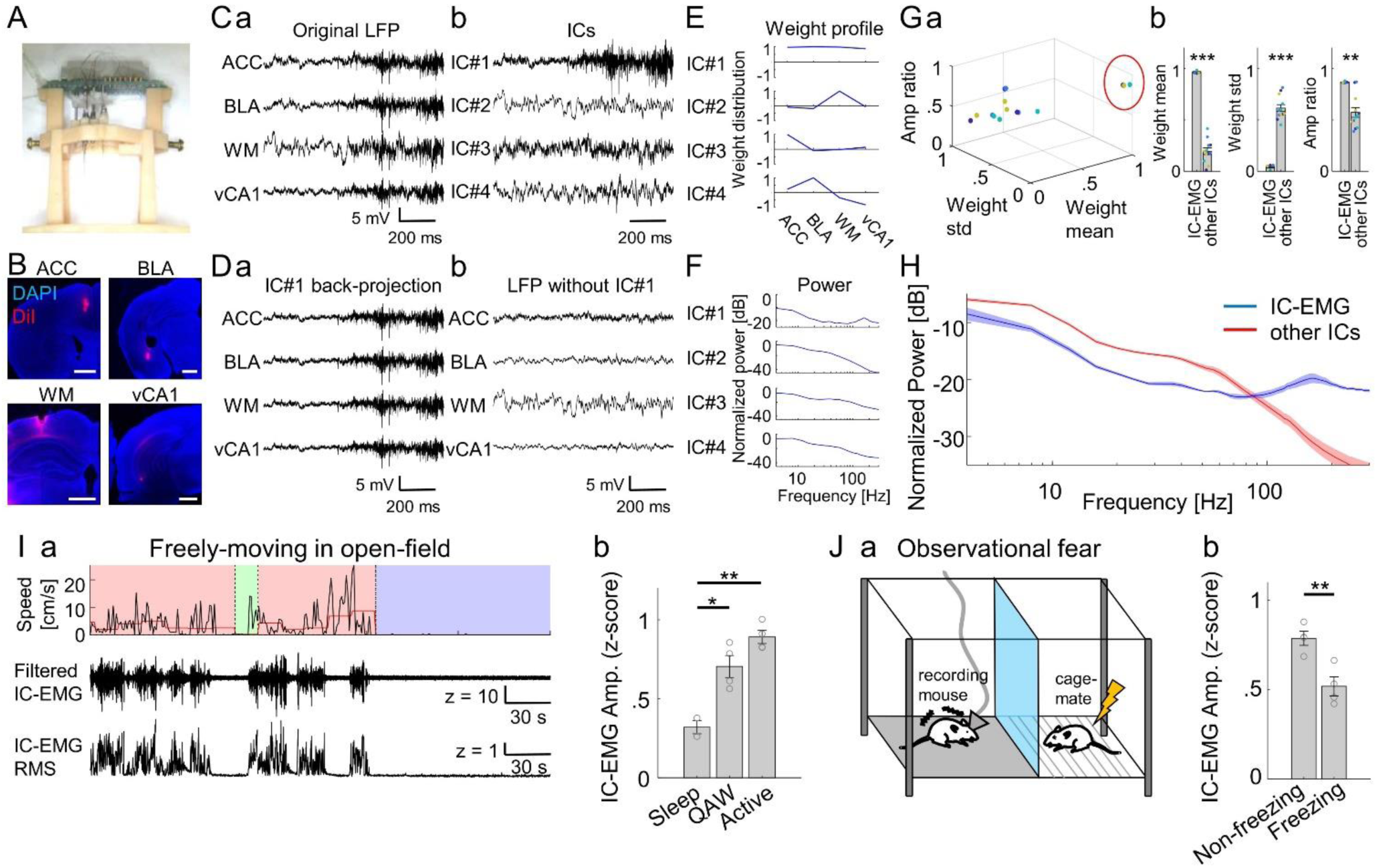
IC-EMG can be obtained from multisite brain-areas recording. (A) Implant device on jig equipped with 4-ch tungsten wire electrodes. (B) The electrode traces. The electrodes were coated with DiI before implantation. Scale bar: 1 mm. (C) Representative LFPs from 4-ch electrodes (a), and their ICs (b). The LFP signals were referenced to the skull screw. (D) The back-projected field potential from IC#1 (a), and the cleaned LFP waveforms obtained by deleting IC#1 LFP from the raw LFP (b). (E) Weight distribution of ICs. (F) Power spectrum density of ICs. (G) (a) Scatter plot (a) and bar plots (b) of IC’s mean weight, std, and amplitude ratio, obtained from four mice. Red circle indicates the uniform-ICs (IC-EMG) in each animal. Averaged parameters were: IC-EMG vs. other ICs: weight mean = 0.96 ± 0.007 vs. 0.20 ± 0.033, t_(14)_ = 13.31, p = 2.45e-9; weight std = 0.045 ± 0.011 vs. 0.62 ± 0.033, t_(14)_ = −9.69, p = 1.37e-7; Amplitude ratio = 0.87 ± 0.005 vs. 0.57 ± 0.047, t_(14)_ = 3.53, p = 0.003, respectively. (H) Average power spectrum density of the IC-EMG (blue) and other ICs (red). (I) IC-EMG from freely moving animals. Speed of animal’s head position (top), 50-500 Hz filtered IC-EMG (middle), and RMS-amplitude of IC-EMG (bottom). (b) Average amplitude of IC-EMG from four mice. Amplitudes between Sleep vs. QAW vs. Active: 0.32 ± 0.042 vs. 0.70 ± 0.070 vs. 0.89 ± 0.042, p = 0.013 (Sleep/QAW) and p = 0.0014 (Sleep/Active). (J) IC-EMG during the observational fear task. (a) Schema of the task, (b) IC-EMG amplitude during animal’s active and freezing states. The z-scored amplitudes during non-freezing and freezing states: 0.79 ± 0.040 vs. 0.52 ± 0.054, t_(6)_ = 4.03, p = 0.007.

## Discussion

Although previous works have shown good performance of ICA in the reduction of EMG-like artifacts from field potential data, evidence of to what extent the separated component and the actual EMG signal correlate was lacking. Thus, no prior attempt was made to use this EMG-like component proactively. The current study has shown that the EMG component obtained from multichannel LFP signals by ICA is highly correlated with actual EMG signals. The IC-EMG was further demonstrated to be useful for measuring animal’s sleep/wake and fear freezing behavior as well as actual EMG. We also showed that EMG component separation could also be performed by ICA with multisite brain-areas LFP recording. Therefore, the current study demonstrated that the EMG signal can be reconstructed from multichannel LFP by ICA without any additional surgery or setup, which can be helpful in the measurement of animal behaviors. In addition, our method allows re-examining animal behaviors in previously-obtained LFP data.

### Advantages of IC-EMG

Although techniques used for EMG signal acquisition are simple, simultaneous recording of EMG with brain activity is not a trivial matter because it requires additional surgeries and setups as we described in the Introduction. In the current study, the total microdrive weight used for silicon probe implant and EMG recording was 6.5 g, while the weight would be reduced to 4.7 g in the case that EMG recording was not needed because the electrical interface board for EMG preamp can be removed. We did not observe abnormal behaviors in the adult mice used in this study, but the increased implant weight could interfere with the behavior of more juvenile animals. Stable EMG recording has also been a challenging problem as the recording wires need to remain attached to the animal despite the mechanical stress caused by muscular movements. In our experiments, three out of twelve mice broke the wires; one mouse broke one side of the EMG wires within a week after the surgery, and another two mice broke one side of the wires while receiving fear conditioning foot shocks. Reconstructing EMG signals from LFP using ICA does not require a wire electrode implant. The ground/reference wire used for the LFP recording connected to the skull screw was fully covered with dental acrylic and did not receive mechanical stress after surgery, ensuring stable recording. Therefore, our IC-EMG method not only reduces the requisite effort of additional surgery and setup preparation for EMG recording but also enables stable simultaneous recording of EMG and LFP with freely moving animals, which allows long-term behavior experiments without the risk of a wire break.

Up to now, several methods have been proposed to score animal behaviors without EMG. Video-based sleep scoring has been shown to have a high correlation with EMG-based scoring ^71, 72^. The video-based methods, however, have a drawback in identifying the fine sleep structures because they take 40 seconds of immobility as a threshold to define sleeping, while short sleep bouts of less than 20 seconds have been reported in sleep-deprived rodents ^73^ and of excessive sleepiness in humans ^74, 75^. Video recordings systems complemented with infrared beam breaking ^76^, piezoelectric signals ^77, 78^, and respiration measurement ^75^ were also proposed; however, these again require additional experimental setups. Sleep has also been often determined based on the oscillatory power of LFP or EEG combined with video-tracking without EMG. In these cases, active awake and REM sleep are identified by prominent theta oscillation (6–9 Hz) while slow-wave sleep is characterized by increased delta oscillation (<4 Hz) ^48, 49, 79^ and by the appearance of sharp-wave-ripple and cortical spindles ^20, 80, 81^. Another study further distinguished sleep and quiet awake states using a criterion that delta power is increased during slow-wave sleep than QAW state ^82^. However, there is still a limitation of this approach in fine state discrimination of immobility states. Although theta is prominent during REM sleep, theta power during QAW can also increase with many factors including respiration ^50^, vestibular stimulation ^51^, sleepiness ^52^, fear ^8, 21, 22^, attention and other cognitive functions ^53–55^. In addition, QAW theta becomes less prominent during waking immobility, eating, grooming, and defecation ^53, 83^. Also, QAW delta power is reported to increase prior to the wake-to-slow-wave-sleep transition onset ^84, 85^. Thus, EEG-based behavior classification has difficulty in fine discrimination of QAW, slow-wave sleep, and REM sleep. IC-EMG provides the benefit of precise animal state discrimination compared to other conventional methods without utilizing EMG information.

While we demonstrated that IC-EMG can be a measure of sleep/wake and fear freezing behavior, we also observed its amplitude was increased during water-licking, food-chewing, and grooming behavior. Thus, IC-EMG has potential to be a measure of various animal behaviors, although further investigation is needed to clarify to what extent animal behavior types can be scored by IC-EMG.

In this study, we obtained EMG signals from LFP data by ICA. The same methodology has potential to be applied to EEG research, considering that ICA has been a well-established method to eliminate EMG artifacts from EEG data. Recently, several source separation techniques, including empirical mode decomposition ^86, 87^, an improved version of ICA-based separation ^88^, and others ^89–92^, have been proposed to have better noise reduction quality for EEG data than conventional ICA with the condition of fewer recording channels. Therefore, utilization of these signal source separation techniques would be possible to improve the quality of EMG-signals isolation from EEG data.

### Signal Source of IC-EMG

We obtained IC-EMG from the multichannel LFP signals, which were characterized to have uniform IC weight distribution. Considering neural activity factors surrounding the electrode should contribute to LFP in a more localized manner ^24, 93^, the signal-source of a such uniform distribution component is supposed to be distal volume conduction ^94^ or a common electrical reference ^34^. However, the volume conduction factor is less likely to be the main factor of IC-EMG. First, IC-EMG has a peak power at 130–140 Hz, while power line conduction should generate a peak at a lower frequency range. Second, such a high-frequency neural event in a brain corresponds to sharp wave ripple ^20^, but it is unlikely to be generated from a uniformly distributed source and its amplitude should be smaller than other slower oscillatory activities. Thus, although both signals were possibly separated by ICA, we assume that the major factor of that uniform component is the electrical reference. In this study, we used the skull screw over the cerebellum as an electrical reference, which can pick up both signals of muscle activities near the screw and cerebellar activity. However, although the precise contribution of cerebellar EEG is unclear, it is reported that overt activity is not seen in a healthy rodent brain ^95^. Together with the high correlation with neck-EMG, we assume the head-neck muscular activities surrounding the electrically referenced skull screw is a major signal source of IC-EMG.

### Factors that may affect IC-EMG identification

In our method, the high-noise level channels identified by eye were removed prior to running ICA. Such noisy signals can be caused by the degradation of the silicon probe electrode that leads to crosstalk between channels ^96^ and changes in the frequency-impedance characteristics ^97, 98^, which have a risk of skewing the time-course of LFP traces. Because ICA is a technique to separate temporally maximal independent waveforms, the skewed LFP can generate additional components for which signal-source actually does not exist. Indeed, although only one IC-EMG component was obtained from each mouse in the current study after the removal of bad channels, we have observed two uniform components correlated with EMG (mean weight = 0.97 and 0.95, mean std = 0.02 and 0.03, Amp ratio = 0.65 and 0.46, correlation coefficient = 0.97 and 0.97, respectively) from one mouse when we did not remove the channels of high noise level before the ICA computation. Thus, including unhealthy electrode site signals in ICA may generate “ghost” components, although EMG signals can still be reconstructed from the LFPs.

### Benefits of using ICA for reducing noise arising from outside-brain reference electrode setting

In this study, ICA was demonstrated that it can separate muscle activities from multichannel LFP data with a skull screw electric reference. Another advantage of applying ICA to LFP data is, as we demonstrated in Fig. 2Bb, it can obtain noise-cleaned LFP data against a brain-external reference.

There have been multiple methods for reducing the noise arising from the distal reference electrode, which is set to avoid the signal distortion caused by placing the reference electrode close to the recording site. Common average reference is the method to generate a more ideal reference by averaging all the recordings on every electrode site to use as a reference ^30^. While the ideal reference can be obtained if the average produces near-zero activity, the average does not often produce zero especially in the case of recording from the local brain area because neural activities at each recording channels are supposed to be correlated each other. Current source density analysis has also been widely used as a reference-independent measure which estimates the strength of extracellular current generators ^31–33^, which is conducted by calculating a second spatial derivative of the recorded potentials between neighboring electrodes in the standard method. However, because the spatial noise is amplified through the differentiation, it often requires spatial filtering ^99–101^ which decrease spatial resolution of LFP signals. In addition, this method is not useful in the case of certain recording electrode setups such as those using tungsten wire electrodes because it requires precise electrode site distance information, nor in the case of multi-brain site recording. The ICA-based method, on the other hand, does not depend on the electrode locations, and subtracting electrical reference activity can be interpreted as re-referencing to a zero-activity location ^34^. Thus, our method to use ICA to LFP data does not only have an advantage in extracting EMG activities without an EMG electrode implant, but also in obtaining noise-reduced LFP signals avoiding the potential signal distortion that is encountered in the case of a within-brain reference electrode placement.

## Conclusion

This study certified EMG signals can be reconstructed from multichannel LFP signals by ICA. Thus, our method provides a physiological measure of animal behavior in LFP data without the extra effort of direct EMG recording. It will be a useful tool in wide-ranged in vivo electrophysiology experiments including investigating neural correlates of animal behavior.

## Acknowledgement

We thank Dr. William Marks for his comments on the manuscript, and all other members of the Kitamura laboratory for their support. This work was supported by grants from the Endowed Scholar Program to T.K., Brain Research Foundation to T.K. (BRFSG-2018-04), Faculty Science and Technology Acquisition and Retention Program to T.K., the Whitehall Foundation to T.K. (2019-05-38), the National Institute of Mental Health to T.K. (R01MH120134 and R01MH125916) and Japan Society for the Promotion of Science to H.O. (201860198 and 202101654). (Corresponding author: Takashi Kitamura.)

## Author Contribution

H.O. and T.K. contributed to the study design. H.O. conducted all experiments and analysis. H.O., J.Y. and T.K. contributed to the interpretation for data analysis. H.O. and T.K. wrote the manuscript. All authors approved the final manuscript.

## Supplementary Figure Legends

**Fig. S1.**
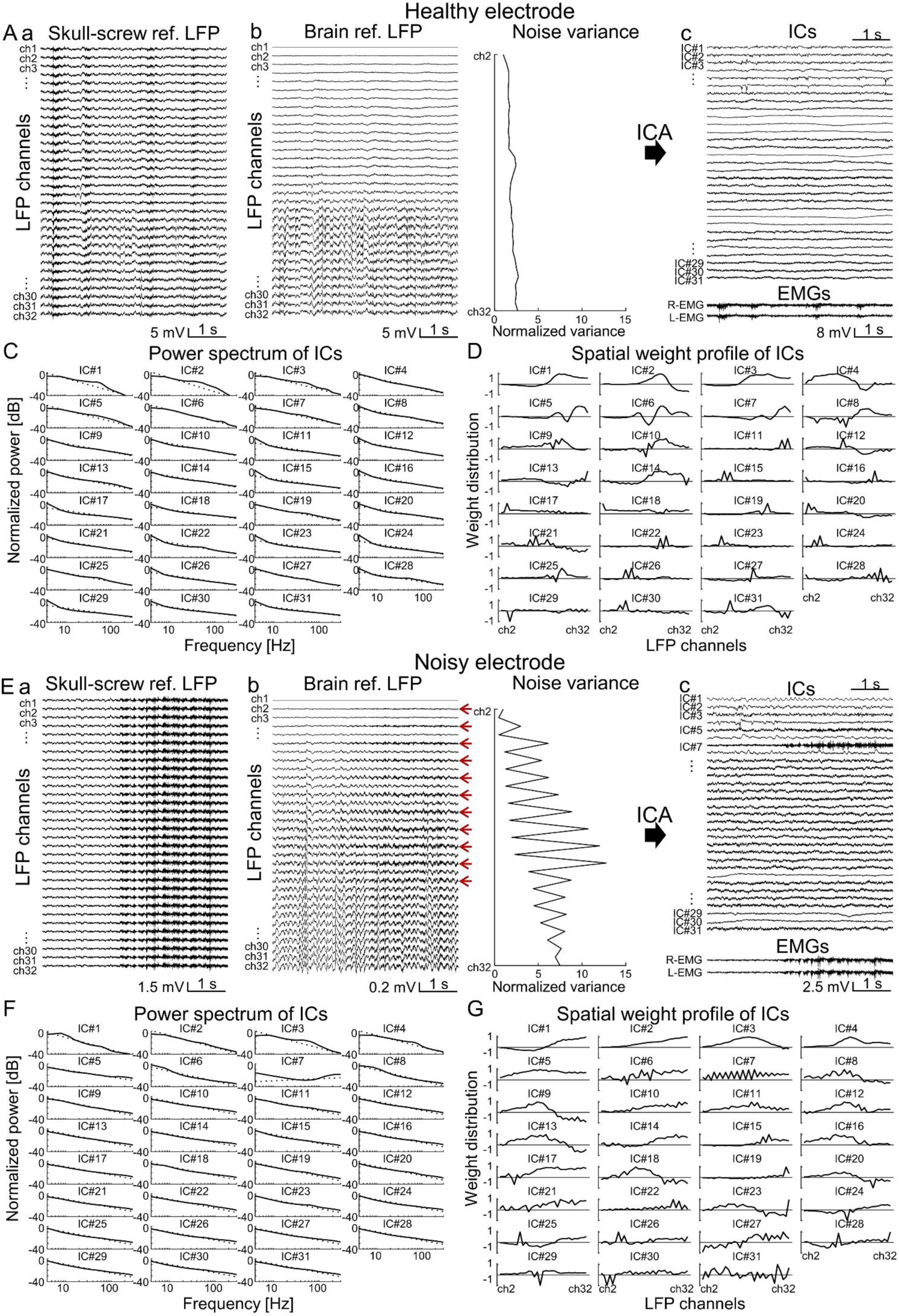
ICs obtained from brain-referenced LFPs, relating to Figs. 2 and 3. (A-C) Results of ICA to LFPs obtained from a healthy electrode. (A) (a) Skull-screw referenced raw LFPs which are same with Fig. 2Aa. (b) Left, brain-referenced LFPs which are calculated by subtracting ch1 activities of the skull-screw referenced LFPs. Right, noise variance of the brain-referenced LFPs, calculated by variance of 150-500 Hz bandpass filtered LFPs followed by normalizing with the variance at ch2. (c) ICs obtained by ICA to the brain-referenced LFPs (top) and raw EMGs (bottom). EMG-like component was not observed in the ICs. Channel#1 of the brain-referenced LFP was not included in ICA calculation (thus 31-ch LFP data was used for ICA) because it only has zero-values. (B) Log-log plots of the normalized power spectrum densities of each IC shown in *Ac*. Black dotted lines indicate the power of aperiodic exponent. (C) The weight distribution of ICs shown in *Ac*. (D-F) Results of ICA to LFPs obtained from a noisy electrode. (D) (a) Skull-screw referenced raw LFPs. (b) Left, because noise-levels are different depending on the channels, noise remains even in brain (ch1)-referenced LFPs. Red arrows indicate noisy channels compared to the neighboring channels. Right, noise variance of the brain-referenced LFPs obtained similarly to *Ab*, showing even number channels are noisier than odd number channels. (c) ICs obtained by ICA to the brain-referenced LFPs (top) and raw EMGs (bottom). IC#5 and IC#7 show EMG-like activities. (E) Log-log plots of the normalized power spectrum densities of ICs shown in *Dc*. (F) The weight distribution of ICs shown in *Dc*.

**Fig. S2.**
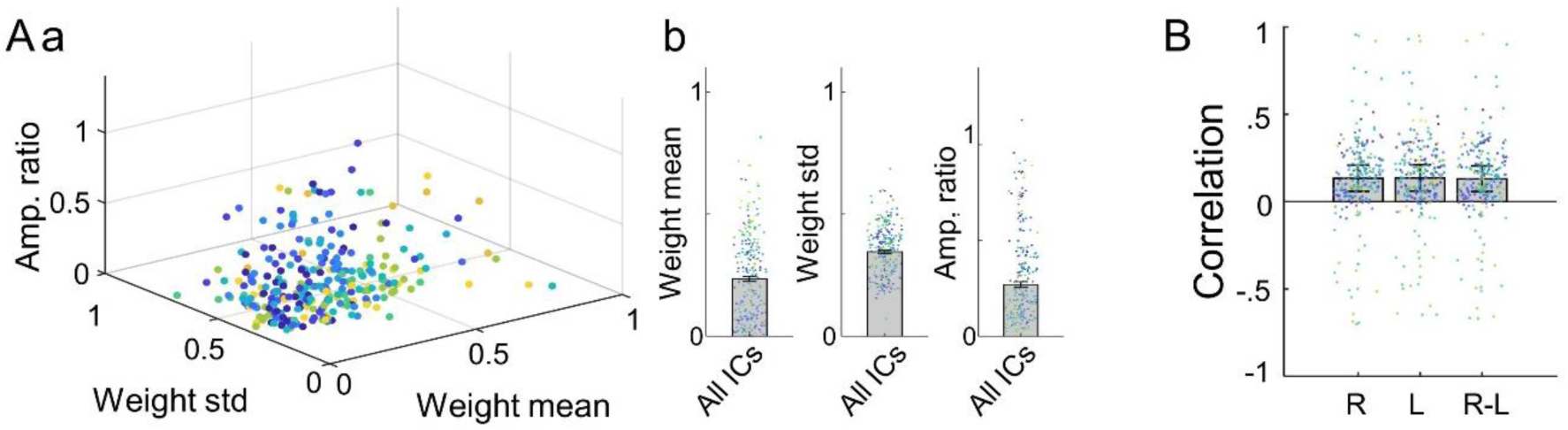
Weight distributions of ICs of brain-referenced LFPs, and correlations with EMG, relating to Figs. 2 and 3. (A) (a) Scatter plot of IC’s mean weight, std, and amplitude, obtained from ICs of the brain-referenced LFPs from ten mice. (b) Averaged parameters of ICs. We could not find clear criteria of EMG-like ICs of the brain-referenced LFPs using these parameters. (B) Average Pearson’s correlation coefficient of ICs (50-500Hz filtered) of brain-referenced LFPs and EMGs (R-EMG, L-EMG, and (R-L)-EMG) from ten mice. Dots of each color indicate different mice. Two out of ten mice had ICs that were highly correlated (>0.9) with EMG, which are obtained from LFPs of noisy electrode recordings.

## Supplement Material

### KEY RESOURCES TABLE

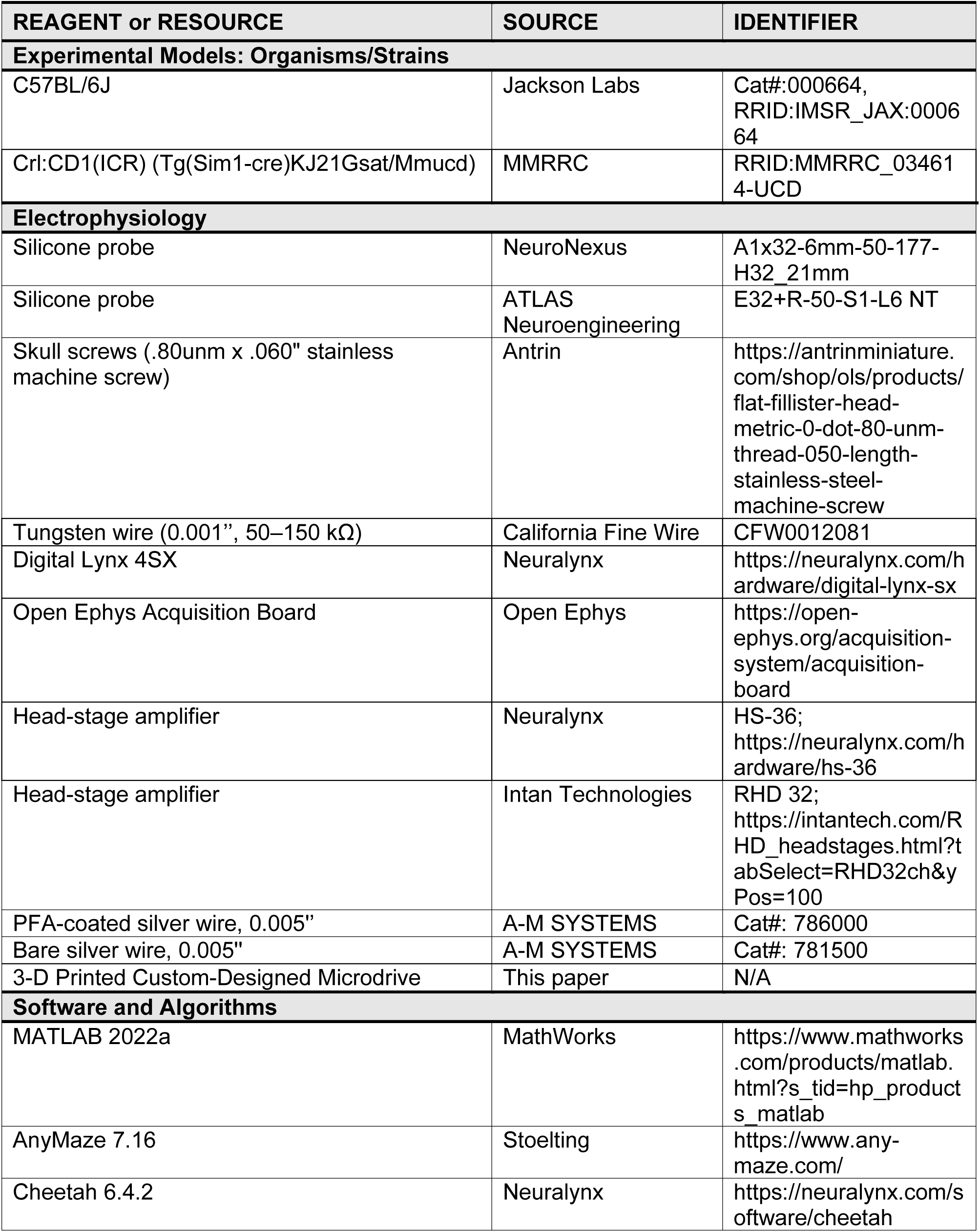
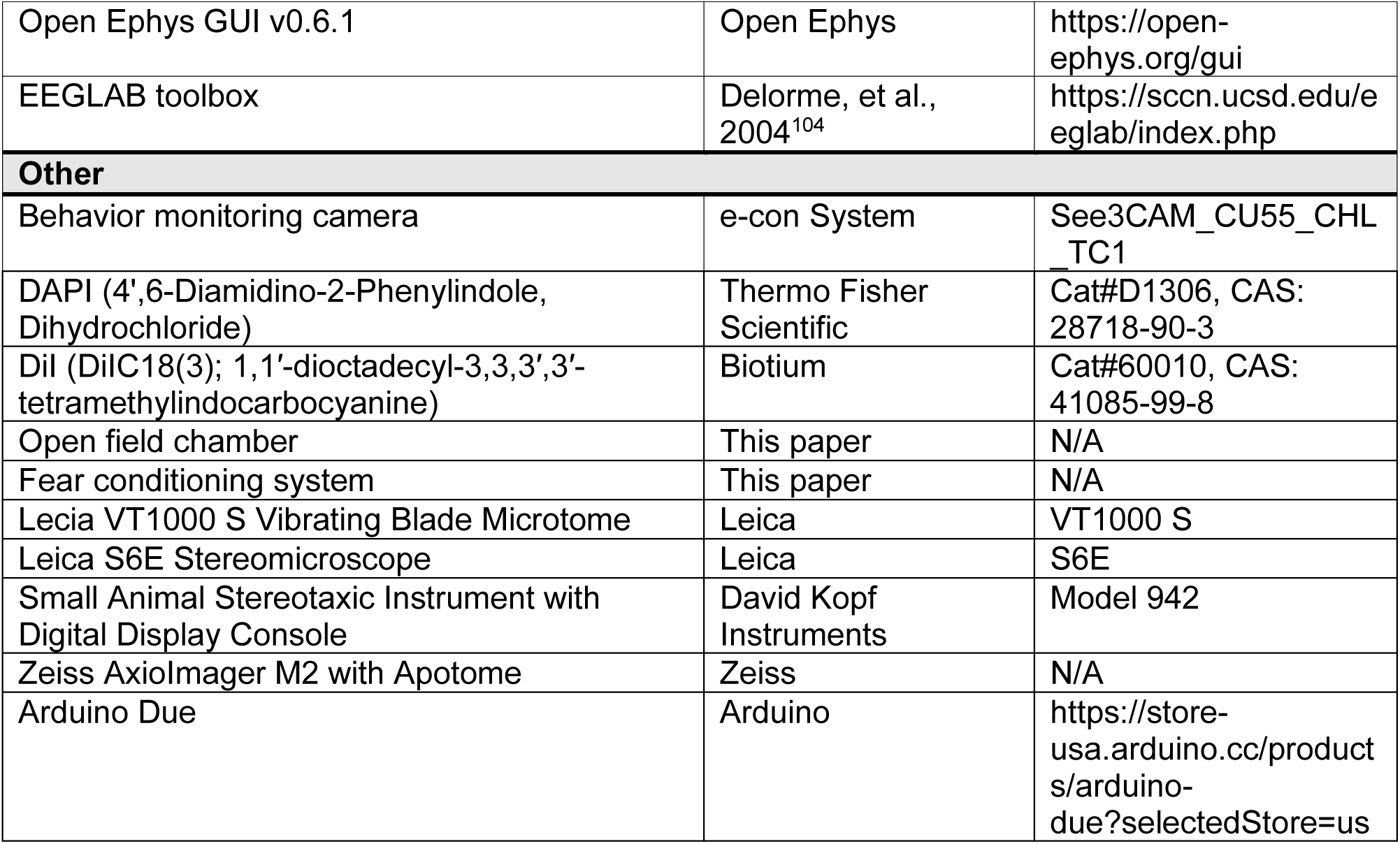

### Lead Contact and Materials Availability

All data are available in the main text or supplementary materials. Requests for resources and additional information should be directed to and will be fulfilled by the Lead Contact, Dr. Takashi Kitamura.

### Method details

#### Animals

All procedures relating to mouse care and experimental treatments conformed to NIH and Institutional guidelines, and were carried out with the approval of the UT Southwestern Institutional Animal Care and Use Committee (IACUC). A total of 16 male mice aged between 2 and 10 months were used. Five C57BL/6J background mice were used for open-field recording. Five Crl:CD1(ICR) background mice crossed to C57BL/6J were used both for open-field and fear conditioning recording. An additional two mice Crl:CD1(ICR) in addition to the four C57BL/6J used above were used to record including REM sleep periods (one mouse was also used for open-field/fear conditioning recording). Another four C57BL/6J background mice were used for open-field and observational fear test with four-channel multisite brain areas tungsten electrode implants, from which data were also used in previous works ^67^. All animals were housed on a 12h/12h light schedule.

#### Electrode implantation surgery

Fabrication of silicon-probe microdrive and implant surgery was based on a previous report ^105^. An implant microdrive equipped with a single-shank 32 channel, 50 μm electrode-site spacing silicon probe (A1×32-6mm-50-177-H32_21mm; NeuroNexus, Ann Arbor, MI, USA; or E32+R-50-S1-L6 NT, ATLAS Neuroengineering, Leuven, Belgium) were prepared for implantation into isoflurane-anesthetized mice. The microdrive was also equipped with a custom-made electric interface board for EMG recording (Fig. 1A), which increased the total implant weight to 6.5 g. In the first step, stainless machine screws (Antrin, Fallbrook, CA, USA) were implanted into the skull to anchor the microdrive. One of these skull screws over the center of a cerebellum was attached with bare silver wire, which serves as an electric ground and reference for the LFP recordings (Fig. 1B). Electric connectivity was checked by measuring that the impedance is less than 20 kΩ at 1 kHz between the ground screw and other skull screws. The silicon probe was then implanted into hippocampal CA1 and dentate gyrus (DG) (AP: −1.80 mm, ML: +1.60–1.70 mm, DV: +2.10– 2.40 mm), then the microdrive was fixed to the skull screws by applying dental acrylic. For the EMG recording, the tips of two perfluoroalkoxy-coated silver wires (A-M Systems, Sequim, WA, USA) were exposed and tied to the left and right dorsal neck muscles (Fig. 1C) ^7, 10^. The skin was then covered with dental acrylic.

Implantation of multiple tungsten wire electrodes was reported previously ^67^. Briefly, four sets of tungsten wire electrodes were mounted on a custom designed 3D printed microdrive (Fig. 5A), which was implanted targeting anterior cingulate cortex (ACC), basolateral amygdala (BLA), white matter above dorsal hippocampal CA1 (WM), and ventral CA1 (vCA1) (ACC: AP: +1.00 mm; ML: −0.30 mm; DV: −1.50 mm, BLA: AP: −1.40 mm; ML: +3.40 mm; DV: −5.30 mm, White matter: AP: −2.40 mm; ML: +2.00 mm; DV: −1.00 mm; vHPC: AP: −3.18 mm; ML: +3.75 mm; DV: −4.75 mm). LFP data were re-referenced offline to a skull screw above the cerebellum. All mice were allowed to recover for at least seven days after surgery. All electrode placements were histologically verified with brain slices stained with 4′,6-diamidino-2-phenylindole (DAPI) (Fig. 1D and 7B). The probe was coated with 1, 1′-dioctadecyl-3, 3, 3′, 3′-tetramethylindocarbocyanine (DiI) before the implantation.

#### In Vivo Electrophysiology and Behavior Experiments

LFP and EMG signals (Fig. 1E) were recorded using Neuralynx (Neuralynx, Bozeman, MT, USA) or Open Ephys (OEPS Tech, Lisbon, Portugal) ^106^ systems. The wide-band signals were obtained from the implanted electrodes and the EMG wires at 20 kHz, then band-pass filtered (0.1–500 Hz) and down-sampled to 2 kHz.

The LFPs and EMGs of the ten animals were recorded while exploring in the open field chamber (70 cm × 38 cm) for 0.5 to 2 hours. Subsequently, five out of the ten animals were recorded during the tone fear conditioning task using a custom-made fear conditioning box ^107^. The box consists of a 30 cm × 25 cm × 25 cm transparent acrylic box surrounded by white plastic boards, and 3 mm diameter stainless steel bars with a 6.66 mm pitch on the floor. Conditioning foot shock (1.2 mA for 20 kΩ impedance) was designed to be delivered to the bars while animal behaviors were monitored with a camera (N660 1080P, NexiGo, Beaverton, Oregon, USA) equipped on the side of the box. Before fear conditioning, all animals were habituated to experimenters for 2 days. On Day1, mice were placed in the fear conditioning box and allowed to explore for 240 s. Then, a 20 s tone (90 dB, 8 kHz) was played from a speaker (ST304, VIP PRO AUDIO, Brooklyn, NY, USA), and a 2-s foot shock was delivered to the mice at the end of the tone, and the animals’ behaviors were subsequently monitored for 240 s. This was repeated two more times before the mice were returned to their home cage. On Day2, the mice were placed in the same box but with a different context in the form of black plastic boards on the floor and the surrounding walls. Similarly to day1, the mice explored the box for 240 s, then 20 s tone delivery and 240 s behavior monitoring were repeated in total three times. LFP and EMG were recorded for the whole period of Day2. The timing of the tone and the shock delivery is controlled by the triggering signals of Arduino Due (A000062). The shocker consists of a portable AC power battery (ZeroKor R350), a step-up transformer (Simran AC-100) and a rectifier (KBP210G, Diodes Incorporated, Plano, Texas, USA) for shock delivery, and bipolar transistors (BC547BBU, onsemi, Phoenix, Arizona, USA) and optocouplers (PC817) for shock-timing control. In addition, one mouse used above and two additional mice were recorded in their home cages (18 × 28 × 12 cm) for 4 to 6 hours to record LFP data including REM sleep periods.

Another four mice were recorded during the observational fear task in the previous study ^67^. Briefly, implanted mice were group housed with their cagemate for one week. On Day1, the implanted mice received one time 0.5 mA, 2-s foot shock (Med Associates, Fairfax, VT, USA). On Day2, the implanted mice and their cagemate were placed in the shock and monitoring chambers, respectively, which were divided by a transparent plexiglass partition. The mice were allowed to explore the chambers for 5 min, then the implanted mice observed the cagemate receiving 0.5 mA, 2-sec foot shocks with an interval of 15 s for a total of 24 shock trials). LFPs were recorded from ACC, BLA, WM, and vCA1 during Day2. Due to excessive line and harmonic noises generated during recording in this shock system, we applied notch filters at 60/120/240 ± 0.1 Hz.

#### Data Analysis

Unless otherwise noted, analysis of electrophysiological data and animal behaviors were performed using custom written scripts in MATLAB (Mathworks, Natick, MA, USA).

#### Independent component analysis of LFPs

To the aim of reconstructing EMG signals without direct EMG recording, we performed ICA on LFP data ^34, 108, 109^. Before running ICA, we removed unhealthy channels from the LFP data with two criteria: (1) its waveform is distinct from the neighboring channel (unfunctional channel), (2) the noise level is larger than the other electrode sites that are observable by eye (degraded channel).

To separate source signals from LFP data, we used the infomax ICA algorithm ^35, 110^ which is implemented in the EEGLAB toolbox (*runica.m*) ^104^. In ICA, the relationship between signal sources and the recorded signals is modeled by

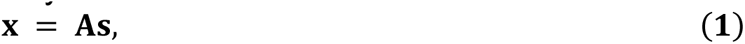

where **x** is the recorded signals, **s** is the source signals called independent components (ICs), and **A** is the unknown mixing matrix which is to be estimated by the ICA algorithms. The source signals are then obtained by the equation

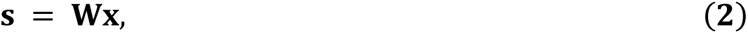

where **W** is computed by the inverse of matrix **A**, which is called unmixing matrix. The number of IC **s** is same with the channel number of the recorded signals **x**. Each column of the mixing matrix **A** describes how the source signals distribute along the original channels, which we called “weight distribution”. The reconstruction from the *n*’th component onto the original data channels is called inverse ICA, which is accomplished by multiplying the *n*’th column of the mixing matrix **A** with *n*’th IC. For comparing IC’s distributions, we normalized their weight distributions so that their maximum absolute is equal to one and their mean value is positive.

Here, we applied ICA to the first 4-min epoch of 32 channel silicon-probe data or previously reported 4 channel tungsten-wire data ^67^ instead of using the full set of data to reduce the calculation cost. The unmixing matrix **W** was calculated through ICA as described above from the 4-min epoch, and then source signals including outside of the 4-min epoch were obtained using equation (2). Although data rank reduction using principal component analysis is often applied before ICA, we did not use it because rank reduction can potentially reduce the quality of subsequent ICA ^111^. While it is known that too many IC separations can cause overfitting, a problem in which IC waveforms are composed of repeated ‘bumps’ that do not appear in the original waveforms ^112, 113^, and indeed we have observed this problem when we have used a 25-μm-spacing 64-ch silicon probe, this overfitting problem was not observed with our recording setups in the current study.

#### Analysis of Power Spectral Density, IC Amplitudes, and IC-EMG/EMG Amplitudes

We used Welch’s method ^114^ with a window length of 0.25 s and an overlap of 0.15 s to estimate the power spectral density of EMG and IC. Aperiodic components of the power spectral density were parameterized ^115^ to help find the spectral peaks. To compare the spectral features of EMG and ICs among different animals, the power spectral densities were normalized by their summation along all frequency ranges.

For evaluating the amplitudes of each IC, we computed how much the ICs account for the original LFP. We calculated the amplitude ratio (Amp. ratio) by the maximum variance of the IC’s back-projected (reconstructed) signals among the channels divided by the variance of LFP at the corresponding channel.

To calculate the time-changing amplitudes of the EEG and the EMG components obtained by ICA from LFP (IC-EMG), we first applied the zero-phased band-pass filter (50–500 Hz) to the original waveforms and then their root-mean-squares (RMS) were obtained with 100-ms time windows. Comparisons of RMS among different animals were conducted with z-scored EMG/IC-EMG data.

#### Behavioral and Statistical Analysis

During the open-field recordings, the animals’ head positions were tracked based on the position of LEDs mounted on the headstage preamplifier. The head position movements were then computed with their average in chunks of every 15 seconds. The video-based mouse behaviors were categorized as Sleep if the head speed was less than 0.2 cm/s lasting more than 40 s ^71^. Animal’s awake states were further categorized as Active if the velocity is greater than 2.0 cm/s and as Quiet Awake (QAW) if the velocity is less than 2.0 cm/s in every 15 s epoch ^82^. In fear conditioning experiments, animals’ freezing behavior was detected by AnyMaze software (Stoelting, Wood Dale, IL, USA). The detection of freezing response during observational fear experiments was carried out using the Video Freeze Fear Conditioning System software (Med Associates) as we previously reported ^67^.

To detect REM sleep periods (see Fig. 6), theta-to-delta ratio^48, 49^ and EMG ^56–60^ amplitude were used in addition to the video-based criteria stated above. First, spectrogram of LFP at CA1 st. lm layer was calculated by continuous wavelet transform with Morse wavelet using *cwt.m* function of Matlab 2022a. LFPs were separated into 10 second epochs to reduce computational cost for the wavelet transform and subsequently concatenated to calculate theta (6–9 Hz)-to-delta (0.1–4 Hz) ratio. Next, video-based classified Sleep state period were re-classified using IC-EMG/EMG amplitude. The 15 seconds chunks are further separated into 5 seconds chunks, then the video-based Sleep was re-classified into QAW if the EMG amplitudes were above z = 1.5 for more than 20% of the chunk duration and otherwise remained into Sleep. Then, Sleep periods were classified into REM if theta-to-delta ratio ≥ 4 ^60^ which continues more than 5 seconds and otherwise into NREM. The short theta-to-delta < 4 periods which are <5 seconds between REM states are also categorized as REM state. Only the mice showing >20 mins sleep durations and showing REM states (six mice) were used for the further analysis. Cohen’s kappa values was calculated by *κ* = (*p*_*o*_ − *p*_*e*_)/(1 − *p*_*e*_) ^64–66^, where p_o_ is observed agreement rate between video-based/IC-EMG-based REM periods vs. EMG-based REM periods, and p_e_ is random agreement rate between them.

We used Student’s t-test after testing normality. Tukey’s honest significance tests were employed for multiple comparisons. p < 0.05 was assumed to be statistically significant. * indicates p < 0.05, ** indicates p < 0.01, and *** indicates p < 0.001. All tests were two-sided. Bar plots and error bars represent means and standard errors.

## Code availability

The custom Matlab code is available at https://github.com/HisayukiOsanai/IC-EMG.git

